# Metabolic inhibition can enable compact physical integration of microbial partners

**DOI:** 10.64898/2026.07.13.738165

**Authors:** Nandakishor Krishnan, József Garay, Ádám Kun, Mark Broom, István Zachar

## Abstract

Microbes often generate by-products that inhibit own or consortial growth. Such imposing constraints can be alleviated by symbiotic partnerships where the partner consumes the limiting factor resulting in mutually beneficial cooperation. Theories of mitochondrial origin envisage such metabolic syntrophy of partners as the initial interaction, often coupled with mechanistic membrane gymnastics to trap the alphaproteobacterial symbiont to ensure vertical inheritance and as a prerequisite or vestibule of endosymbiosis. Yet we do not know how this happened. Here, using a mathematical model, we investigate how such metabolic self-inhibition enables and facilitates the evolution of compact physical integration between syntrophic partners. We show that when one species produces a metabolite that is self-inhibitory, selection favors increasingly tight spatial association of a detoxifying partner to mitigate local accumulation of the self-limiting product. Under a broad parameter range, this process drives the emergence of surface-associated configurations promoting morphological adaptations such as membrane protrusions or invaginations that increase inter-partner contact area. Our results prove that such costly host membrane contortions can evolve as a means to improve benefits of inhibition reduction by protective syntrophic ectosymbionts. Additionally, a unilateral syntrophic interaction between parties is sufficient for the transition; a mutualistic (or bilateral) one is not necessary. Our model establishes a direct evolutionary link between metabolic coupling and physical integration, suggesting that the need to alleviate (self-generated) inhibition can be a primary driver of ectosymbiotic attachment and its transition toward more intimate associations. Our findings provide a general theoretical framework for understanding how microbial consortia evolve structural complexity from initially loose metabolic interactions, with implications for the origins of stable symbioses and the early steps toward cellular integration.

## Background

Syntrophy is an extremely common interaction among microbes in all three domains: bacteria, archaea and eukaryotes; and it is assumed to be a major driver of symbiotic integration of microbial species. Syntrophy, in its broadest definition, is mutualistic metabolism [1] that covers a wide array of metabolite-mediated interactions between microbes, ranging from cross-feeding [2] to joint catabolism [3,4]. For example, methanotrophic archaea and sulfate-reducing bacteria work together to perform anaerobic oxidation of methane [5]. Syntrophy can involve the sharing of nutrients or electrons, unilaterally or mutually, for which examples abound (see, e.g., [6,7]). One particularly important interaction type is the consumption of inhibitive or downright toxic by-products of a species by the partner [8,9] (also modelled extensively [7,10,11]). Such detoxifying behavior is indirectly mutual (and henceforth cooperative) as the producing party feeds the consumer, eliminating the accumulation of the inhibitive product in the local neighborhood. Often, but not necessarily, the consumer generates by-products that are further beneficial to the producer species.

Though syntrophic interactions are mostly considered to be obligate and some are even species-specific [12–14], we believe this is an evolved feature, and multispecies consortia generally do not rely on exact taxonomic combinations [15]. Moreover, multispecies partnerships could not be reliably maintained over generations without effective means of consortial reproduction (something that evolved only in proper multicellular organisms and in endosymbioses), but rather functional recruitment of relevant phenotypes from the environment [16,17]. Accordingly, the integration and maintenance of multispecies genetic information of the different strains pose a problem, likely limiting their coevolutionary potential. This problem is likely more acute in environments that are less stable, with many species in flux compared to the hypothetical cases where only two cooperative species occupy a habitat. Obligately symbiotic species nevertheless exist, e.g., the recently discovered *Prometheoarchaeum* [14,18]. However, how they evolved and ensure (if they can) prolonged presence of their obligate partners is yet unknown. While endosymbiotic capture enables direct vertical inheritance (though not necessarily, think of the reuptake of symbionts per generation in many lineages), it is only available for eukaryotes, as far as we know for now. Yet an effective method to increase co- occurrence of relevant strains, potentially available for prokaryotes too, is the surface-bounding of the partner species, as many ciliate hosts attest [19,20] (for prokaryotic examples, see [6]), which could be a beneficial adaptation when partners exchange metabolites.

As a matter of fact, we do not even know how microbial endosymbioses form. We only have very limited knowledge on how some modern ectosymbioses function and how they were shaped metabolically and genetically [13,21–23] and minimal information on their evolutionary origins. Regarding the mechanism of partner capture, with the exclusion of phagocytosis (unknown among archaea so far), some theories (older and newer [18,24–27]) assume host-initiated inward or outward membrane folds (invaginations or protrusions). However, no example or evidence is known for any such successful capture yet. Nevertheless, microbial consortia do exist in numbers and generally involve some metabolite exchange between partners, presumably dependent on contact area size.

Surface contact interactions have two trivial benefits: close contact not only maximizes the surface area over which metabolites are directly exchanged (uni- or bidirectionally) but also allows for the privatization of these metabolites against third parties in the environment. While metabolic interactions without surface bounding can be beneficial for the parties, they are limited by the available area of direct secretion (by host) and uptake (by symbiont) and are prone to be exploited by selfish third parties or hindered by dilution of the products. In addition, surface bounding of syntrophic symbionts helps in consuming and reducing the local concentration of the host metabolite, and this elimination is beneficial to the host if the metabolite is toxic or self-inhibitory to them [28]. By maintaining surface-bound symbionts, a host that does not depend on the symbionts for metabolites receives an indirect benefit in the form of reduction in self-inhibition. Hence, it is reasonable to assume that to further increase the benefit of the interaction, parties have to increase their joint contact area. Among metabolically coupled, cooperative microbes, the contact surface area seems to be important, that also turned out to be a crucial determinant of advanced physical integration according to models [28]. Arguably, this also extends to endosymbiotic integration (Fig. 1) and to the origin of mitochondria according to some proposed theories [18,25], though the benefit of increased surface contact was never tested.

**Fig. 1.**
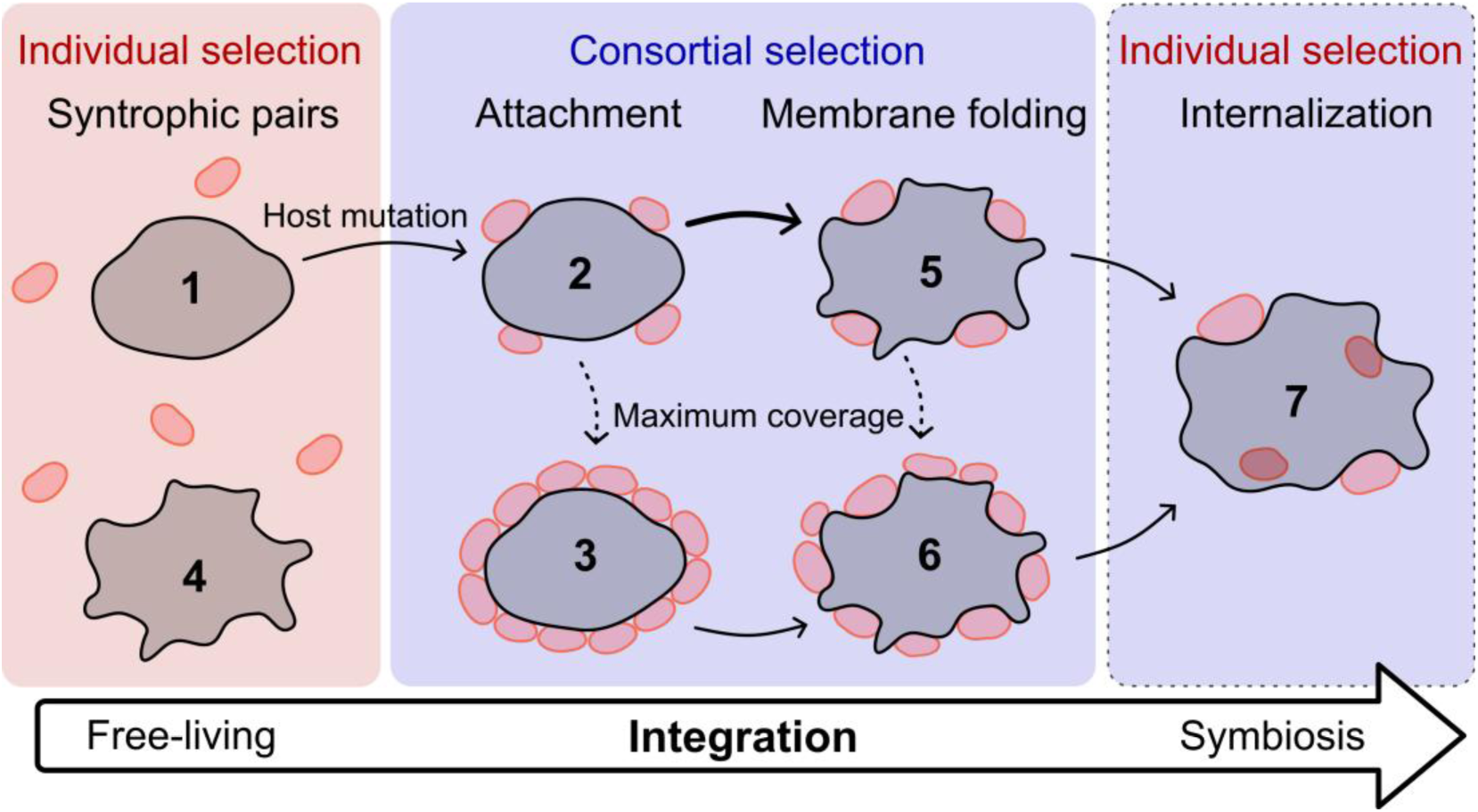
Progressive symbiotic integration. Hypothetical pathway to achieve physically integrated associations and ultimately endosymbiosis from syntrophic, free-living hosts and symbionts. Hosts mutate (denoted using solid black arrows) to form potentially beneficial adaptations leading to better-integrated associations with symbionts. Hosts could also end up with maximum surface occupancy, which is treated as a special case of surface attachment and not a mutation (denoted using dashed black arrows). In this paper, we focus on the evolution of membrane folding as a feature with better surface coverage (compared to typical ectosymbiotic attachment) to reduce self-inhibition to hosts without fully compromising their food influx. We introduce our model by considering an already stable ectosymbiotic consortium where attachment has already evolved and compare two cases: 2 against 5 (see Methods 2) and later generalize it to include other possibilities of interactions (see Methods 3).

The benefit can be direct availability of food for the host (e.g., in case of cross-feeding) or to counter growth inhibition due to harmful factors produced by the host, or both. Here, contact is crucial as the partners are adjacent to each other and molecule transfer between them is directly enabled [29]. Therefore, ideally, metabolite-mediated ectosymbioses should tend to maximize partner proximity by, e.g., developing full coverage of the host surface by the symbionts. However, biomembranes are usually used for food uptake, energy production and other functions that are indeed hindered when covered [30], especially if assuming an autotrophic host (even photosynthetic) where growth is limited by metabolite uptake. Therefore, increasing the ectosymbiont coverage is expected to yield shrinking benefits and, above a threshold, constrained food consumption of the host occupied by too many ectosymbionts would likely render this association unfeasible. Thus, even though the benefit of the interaction depends on contact surface area, the symbiont coverage cannot be increased any further without risking suffocation and the host cannot further increase its benefit. Given this trade-off between inhibition reduction and food uptake reduction based on the contact surface, a different mechanism is required where the host can reduce exposure to toxic metabolites while maintaining some symbiont-free surface so that food influx can still happen. For instance, membrane invaginations, protrusions or appendages can potentially further increase the contact surface with surface-bound partners, without fully compromising surface availability for substrate intake (contrary to full coverage ectosymbiosis).

Here we design a mathematical model (based on previous works) to investigate the feasibility of membrane invaginations or external protrusions as means of increasing the contact interface while accommodating a metabolically active cooperative partner. Using our model, we analyze the hypothesis that forming such membrane contortions could be beneficial for a syntrophic pair and selected for against typical ectosymbiosis. We assume a unilateral metabolic interaction (Fig. 2) rather than a heterotrophic lifestyle (or phagotrophy) of the host and analyze the benefits and costs of contact that can potentially stimulate evolution of mechanisms enabling multi-species microbial integration.

**Fig. 2.**
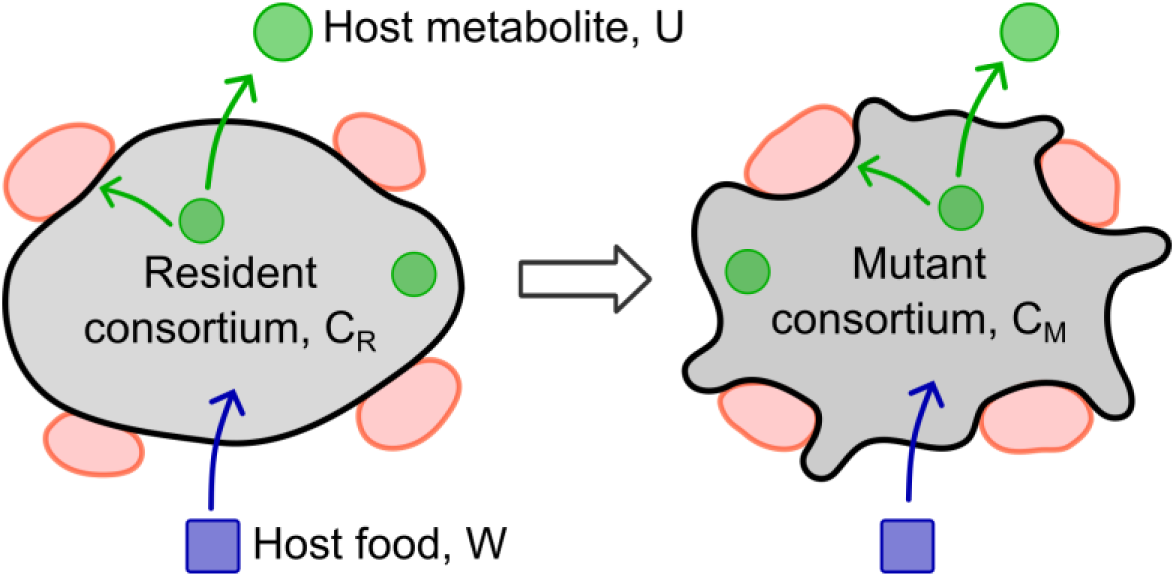
Syntrophically coupled ectosymbiotic consortia. The hosts feed on a substrate W and produce a by-product U that is toxic to them. The host’s inhibitory metabolic product is directly consumed by its ectosymbionts, and excess diffuses into the external environment. The mutant hosts (C_M_) can bend their membrane inward to form invaginations or outward to form external protrusions to increase contact surface with their ectosymbionts compared to their resident counterpart (C_R_). Increased contact increases the beneficial effect (reduced self-inhibition due to reduced exposure) from the ectosymbionts, whereas exposing more unoccupied surface improves food consumption with the additional risk of exposing the host to external inhibitory metabolite bulk. The mutants suffer additional structural and energetic costs to form and maintain membrane contortions, which are expressed in terms of subsistence and replication costs. We investigate the conditions for the evolution of membrane folding in a population of resident ectosymbiotic consortium. The ectosymbiont count and the host membrane size are arbitrary in the illustration. For comparison, the cell volumes and membrane sizes of the two phenotypes are identical. For more details, see Methods 2.

## Results

We test our hypothesis using a model involving hosts and symbionts in unilateral syntrophy, where host growth is limited by a metabolic stress. For simplicity, we first assume the existence of a stable host-symbiont syntrophic pair already engaged in ectosymbiosis, i.e., physical attachment of symbionts on host surface. Next, we analyze the possible invasion of a mutant host phenotype additionally capable of bending its plasma membrane and enfolding its ectosymbionts into it (Fig. 2). For the time being, we consider only two host phenotypes (both ectosymbiotic), later to be generalized to include more cases as in Fig. 1. Membrane bending (or folding) results in increased contact surface between the host and its ectosymbionts, effectively mitigating the effects of metabolically induced self-inhibition (by reducing exposure) to the host compared to that in resident ectosymbiosis. However, the mutant’s membrane bending ability increases costs of subsistence and replication to the host, reflecting the immediate energetic cost of expressing, transferring, installing and maintaining associated proteins. Additionally, to model the worst-case scenario, attached symbionts also hinder the host’s food consumption, impacting its growth and fecundity dependent on the free external surface. The biophysical mechanism of shielding of the host from the self-produced metabolite and reduction in its local concentration while in ectosymbiosis is analogous to diffusive boundary layers common in aqueous microbial contacts. We analyze when and how the costly, membrane-bending mutant phenotype of the host can invade the resident system.

We first analyze the existence and stability of the unique steady state of the resident system comprising of resident ectosymbiotic consortium and concentration of its metabolite (see Methods 1). The stability of the unique interior fixed point of the corresponding dynamical system ensures the ecological stability of the resident host-ectosymbiont consortium. Self-inhibition, modelled based on metabolite-based microbial models [31], demonstrates to be suitable for stabilizing the syntrophic pair. The resident ectosymbiotic consortium is unconditionally stabilized by unilateral syntrophy and metabolite-based self-inhibition (details in Methods 1, also see [28]).

Now, when a membrane-folding mutant of the host is introduced into the system of the stable resident syntrophic pair, the extended dynamics with self-inhibition could evolve either to a resident-only or mutant-only steady state (see Fig. 3; details in Methods 2). However, the costly membrane-folding mutation can never invade a resident system of ectosymbiotic consortium stabilized only by intraspecies competition (given there are no additional benefits to the mutant in this case). Density-dependent, competitive interaction does not provide the required stability to the system, entailing metabolite-dependent population saturation (for details, see Appendix E in Additional file 1). Hence, membrane-folding does not emerge spontaneously without the additional presence of metabolic stress; this suggests the necessary role of other factors, such as self-inhibition, in the evolution toward advancing multi-species microbial integration.

**Fig. 3.**
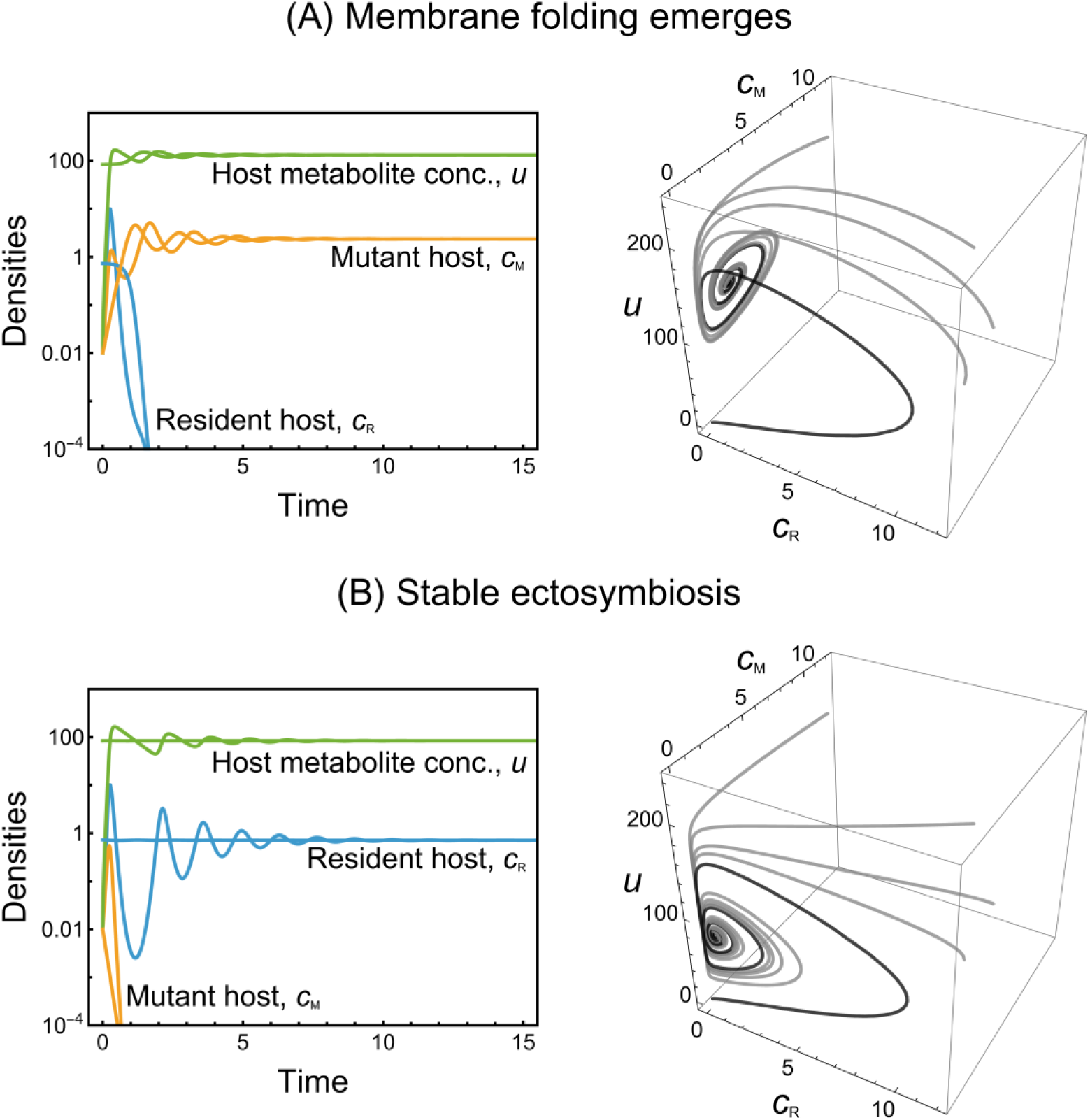
Stability of the resident ectosymbiotic consortium and the membrane-folding mutant host. Time series plots and phase portraits of the dimorphic system are visualized. In the time series plots (left column), initial conditions are arbitrary points close to the trivial and resident fixed points (illustrating local stability). In the phase portraits (right column), the solution for an arbitrary point close to the trivial fixed point as the initial condition is colored black; all other trajectories correspond to arbitrary initial conditions (illustrating global stability). c_R_(t) and c_M_(t) denote time-variant population densities of resident and mutant hosts, and u(t) is the total concentration of the inhibitive host metabolite in the habitat. The parameter values used here are in Table 2. (**A)** The dynamics evolves to stabilize at the steady state 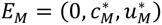, where only the mutant membrane-bending host is stable, for contact area s_M_ = 0.4. (**B)** The dynamics evolves to stabilize at the steady state 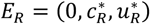, where only the resident host is stable, for contact area s_M_ = 0.25. For more details on the model, see Methods 2. Higher value of contact surface s_M_ = 0.4 by membrane folding as opposed to a value of 0.25 shifts the stability of the system from the resident equilibrium to the mutant equilibrium. Better contact surface area of the membrane-folding mutant is beneficial in reducing the effects of self-inhibition; if these advantages outweigh the cost of reduced intrinsic growth rate, membrane folding can emerge and stabilize.

The extended dynamics with self-inhibition (Methods 2) can only have two ecologically feasible, non-trivial, monomorphic fixed points existing conditionally; a mixed equilibrium (interior fixed point) of resident and mutant host (both with ectosymbionts) is not feasible. Table 1 contains the existence and stability conditions of the fixed points. The existence conditions suggest that the intrinsic growth rate of the hosts should be less than the maximal self-inhibition for the population densities to be bounded and denying population surge. Additionally, the ratio of rate of release (by the host in consortium) to rate of indirect consumption (by ectosymbionts) of the host metabolite should be high. The ratio should be larger than a threshold determined by the host’s growth rate and growth inhibition factors (see Table 1 and Methods 2). We analyze the stability conditions of the mutant-only fixed point in the extended dynamics, i.e., the mutant invasion condition for the membrane-folding phenotype of the host to successfully invade and replace the resident (for examples where similar methodology was used, see [32–36]). Effectively, our model integrates population and metabolite dynamics to analyze specific conditions that could favor invasion of membrane folding in syntrophic microbes.

**Table 1.** The fixed points in the dimorphic system and their existence and stability conditions. The dimorphic system is described in Methods 2 (also see Fig. 3 and Table 2). The trivial fixed point (0, 0, 0) is always unstable.

| Fixed point | Existence condition | Stability condition |
| --- | --- | --- |
| $E_R = (c_R^*, 0, u_R^*)$<br>(resident-only) | $r_R < \frac{1}{b} \ \& \ \frac{\kappa_R}{\xi_R} > \frac{r_R a_R}{1 - r_R b}$ | $\frac{r_M a_M}{1 - r_M b} < \frac{r_R a_R}{1 - r_R b} \ (\Rightarrow u_M^* < u_R^*)$ |
| $E_M = (0, c_M^*, u_M^*)$<br>(mutant-only) | $r_M < \frac{1}{b} \ \& \ \frac{\kappa_M}{\xi_M} > \frac{r_M a_M}{1 - r_M b}$ | $\frac{r_M a_M}{1 - r_M b} > \frac{r_R a_R}{1 - r_R b} \ (\Rightarrow u_M^* > u_R^*)$ |

**Table 2.** Model parameters. The parameter values are arbitrarily chosen for numerical analyses and are compatible with the model descriptions and conditions. For details, see Additional file 1.

| Parameter | Meaning | Arbitrary parameter value |
| --- | --- | --- |
| $w$ | Constant concentration of food source available to the hosts | 500 |
| $\Omega$ | Volume of the habitat or medium | 10 |
| $k_R$ | Consumption rate of the resident hosts | 0.4 |
| $k_M$ | Consumption rate of the mutant membrane-bending hosts | 0.32 |
| $l_R$ | Cost of living of resident hosts | 5 |
| $l_M$ | Cost of living of mutant membrane-bending hosts | 6 |
| $z_R$ | Cost of reproduction of resident hosts | 4 |
| $z_M$ | Cost of reproduction of mutant membrane-bending hosts | 5 |
| $d_R$ | Intrinsic death rate of resident hosts | 0.33 |
| $d_M$ | Intrinsic death rate of mutant membrane-bending hosts | 0.4 |
| $e_R$ | Consumption rate of ectosymbionts on the resident hosts | 0.2 |
| $e_M$ | Consumption rate of ectosymbionts on the mutant hosts | 0.3 |
| $b$ | Reciprocal of maximal growth inhibition to hosts | 0.01 |
| $a$ | Coefficient of growth inhibition to free-living hosts | 1 |
| $a_R$ | Coefficient of growth inhibition to resident ectosymbiotic hosts | 1.67 |
| $a_M$ | Coefficient of growth inhibition to mutant ectosymbiotic hosts | 2, 5 |
| $n$ | Average ectosymbiont count in a consortium | 2 |
| $s_R$ | Surface impact on growth inhibition per ectosymbiont on the resident host | 0.2 |
| $s_M$ | Surface impact on growth inhibition per ectosymbiont on the mutant host | 0.25, 0.4 |
| $\delta$ | Decay coefficient of the host metabolite | 1 |

The fixed-point stability analysis of the extended dimorphic system with both host phenotypes (Methods 2) shows that the mutant invasion condition condenses to a sufficiently high enough inhibition reduction that outweighs the costs incurred (see Fig. 4). Specifically, lower levels of self-inhibition to the mutant host compared to the resident could facilitate the stability of the mutant-only equilibrium. Contextually, this can be achieved only if the mutant membrane-bending host has larger contact surface area with their ectosymbionts compared to that of the residents (which is what we have modelled). Note that larger contact surface is only a necessity under the constraints of our model; there are other ways to reduce self-inhibition such as the symbiont evolving to be syntrophically more efficient in diminishing the inhibitory product. For instance, syntrophic associations where the consuming partner becomes more efficient at removing the host’s inhibitory product is positively selected for (e.g., [3,37]). Similarly, motility enables the host to evade the inhibitory metabolite and reduce the toxic exposure. In fact, host motility may work against improving close contact, unless the symbiont is in obligate physical attachment (either mutualistic or parasitic).

**Fig. 4.**
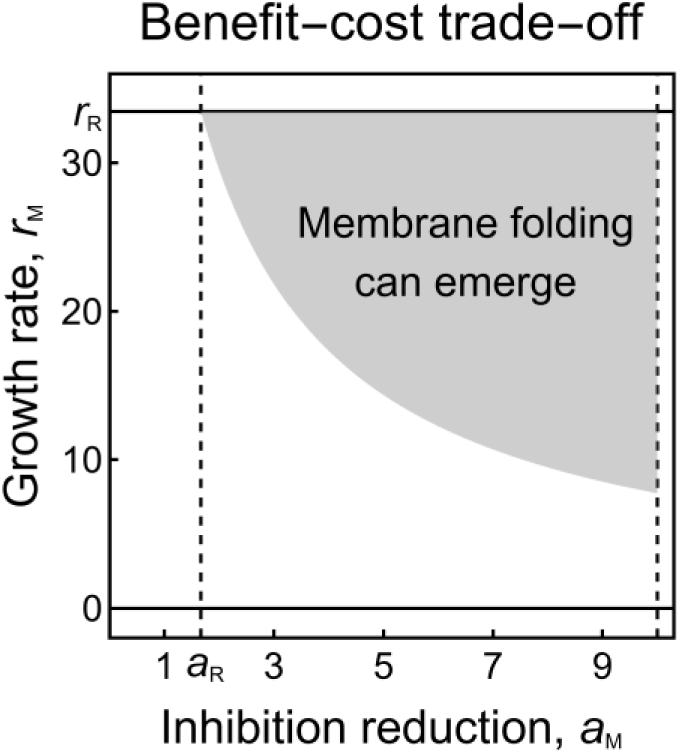
Contact-dependent trade-off to the mutants. The mutant invasion condition depends on a non-linear trade-off between inhibition reduction (benefit) and reduced growth rate (cost) to the mutants. Membrane folding evolves in hosts if it provides a high enough inhibition reduction that can sufficiently compensate for the costs incurred in terms of reduced growth rate. Here, r_R_ and a_R_ denote the growth rate and inhibition reduction factor of resident hosts and are respectively the upper bound of r_M_ and lower bound of a_M_ (upper bound of a_M_ is based on existence condition in Table 1). Higher values of growth inhibition factor (a_M_ ) correspond to lower levels of self-inhibition.

Though the mutant pays extra (area-dependent) costs, which are reflected in its reduced food intake and intrinsic growth rate, the benefits of inhibition reduction per surface area could outweigh it. If it does, the costly mutant stabilizes (Fig. 3(A)). Note that both cost and benefit depend on the common contact surface area, and the trade-off is not linear (Fig. 4). The invasion condition, fundamentally based on the common surface area, manifests a balance of benefits and costs to the mutant, where it can be tipped to receive a net benefit by increasing the contact surface via membrane folding. Thus, the membrane-folding host phenotype with higher contact surface can stabilize and dominate purely based on its advantage of inhibition reduction. Contrary to full-scale ectosymbiosis where the host surface is fully occupied by the symbionts, membrane folds can effectively increase contact surface between the partners *without* fully eliminating host’s free surface (i.e., without compromising processes of food uptake, signaling, interactions, etc.) and with only costs accounting for membrane enfolding. The host’s incentive for symbiont capture in membrane folds relies on the balance between reducing risk of exposure to inhibitive or even toxic metabolite and maintaining some free external membrane surface for food uptake. The trade-off exists between inhibition reduction and metabolite influx at the contact surface. Membrane-folding can achieve more contact surface with the same amount of ectosymbionts, without the need for sacrificing more of active host surface. However, if membrane folding is too expensive for the host (effectively, growth rate is low), they are better off without enfolding the symbionts and they remain in the resident ectosymbiotic form. In other words, if the mutant invasion condition (based on the trade-off) is not satisfied, the mutant perishes and the resident ectosymbiotic consortium remains stable (Fig. 3(B)).

The mutant invasion condition also corresponds to a stable mutant host phenotype capable of tolerating higher concentration of the host metabolite at the respective fixed point (see Table 1 and Fig. 3). Improved tolerance to self-produced inhibitory (or even toxic) metabolites is sufficient to facilitate physical integration in syntrophic, ectosymbiotic partners, given the mechanism to fold membrane is present. Physical integration within membrane folds therefore could evolve in microbes as a mechanism to mitigate the necessary side-effect of their autotrophic growth: metabolic self-inhibition.

Though we have observed the necessity of better physical integration via a mechanism to increase host-symbiont contact surface to counter inhibition by self-produced metabolite, the question remains whether non-self-produced metabolites could facilitate the same transition or not. We demonstrate (in Appendix F in Additional file 1) that even when a metabolite appearing (i.e., produced neither by host nor by symbiont) and periodically varying in the environment is considered, the membrane-bending phenotype can exclusively survive with a sufficiently high contact surface, even against the odds of a higher level of toxicity. Different toxicity levels and ratios of the growth inhibition factors can lead to the existence and collapse of either the resident or the mutant phenotype (Fig. F1 in Additional file 1). Hence, physical integration to improve tolerance against toxic metabolites in the environment can evolve even if such metabolites are externally produced (e.g., by other species) and their concentration is periodically fluctuating, making our result even more robust.

Note that our analysis so far has been based on the dimorphic system with the resident and mutant ectosymbioses without full surface coverage. However, free-living hosts and ectosymbioses with maximum coverage can also possibly exist as special, extreme cases of attachment (though not as a new mutable trait). Now if we assume the attachment mutation in a purely self-inhibitive environment, we have three different possibilities of association (cases 1 to 3 in Fig. 1; also see Methods 3). Similarly, once attachment trait has already evolved, considering membrane folding mutants in the same environment gives separate ecological dynamics (cases 4 to 6 along with stable attachment). In either of the two ecological dynamics (or subsystems), the stability of the dominating type is based on the trade-off between intrinsic growth rate and inhibition reduction. In the system with the membrane folding trait, the cost of limited food intake due to maximum surface occupancy in case 6 in Fig. 1 is likely to outweigh their benefits of least exposure and reduced inhibition (similarly for case 3 when considering attachment mutation dynamics). In case 4, the high levels of inhibition could easily outweigh the advantage of unconstrained food influx (similarly for case 1 when considering attachment mutation dynamics). Therefore, cases 1, 3, 4 and 6 in Fig. 1 are less likely to stabilize in their own respective subsystems based on biophysically plausible hierarchies of growth rates and inhibition factors of the different types (see Methods 3). However, case 5 can be easily stabilized with only a sufficiently high (but not maximum) surface coverage. Moreover, in a purely competitive environment without metabolic inhibition, the free-living (non-ectosymbiotic) hosts with the highest intrinsic growth rate among all the competitor types dominate and cannot be replaced. This result suggests that an additional metabolic stress from the environment promotes the emergence of a host phenotype enabling improved contact with their symbionts. Thus, improved microbial integration can be achieved by the increased contact surface in an attempt to counter the impact of self-inhibition. Even if the phenotypes with higher contact surface simultaneously suffer additional costs in terms of the life history parameters and constrained consumption, enhanced physical integration (via mechanisms like membrane folding) could still evolve in an environment of self-inhibition and unilateral syntrophy (see Methods 3 and Fig. 5).

**Fig. 5.**
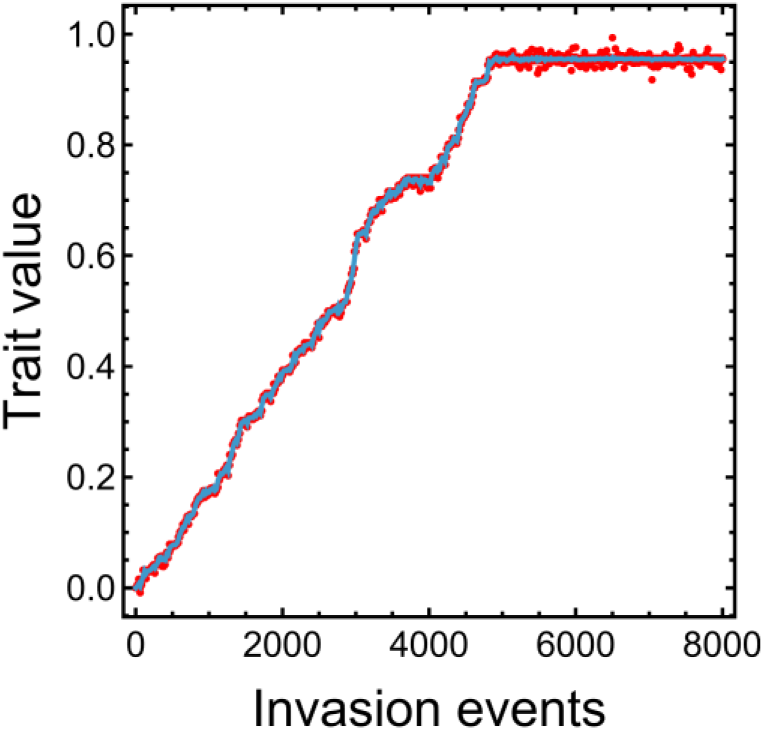
Trait invasion analysis. Variable trait in invasion analysis is driven to the optimal value satisfying the invasion condition. Higher trait values correspond to improved contact and better integration. In an environment of metabolic stress, ectosymbiotic partners tend to develop mechanisms that increase inter-partner contact.

## Discussion

Syntrophic hypotheses of symbiogenetic origins of eukaryotes envisage metabolic exchange as the initial interaction of the partners, eventually leading to a physically associated symbiosis. Presuming that evolution toward an endosymbiotic consortium from a free-living syntrophic pair involved several intermediary transitions, each step would then be due to successful invasion by a beneficial mutation capable of improving the extent of physical association between the host-symbiont pair (e.g., Fig. 1). The mutant invasion results in the formation of a symbiotic consortium, which is a higher level of organization. In this paper, we focused on the emergence of a membrane-folding mutant host capable of enfolding its plasma membrane (either as invaginations or protrusions) to accommodate and hold the ectosymbionts. Here, membrane restructuring serves as an example of mechanism to improve inter-partner integration. The model is based on previous works on:

- Evolution of ectosymbiosis based on symbiont- or host-evolved features [28,38], exploring the possibility of (obligate) attachment between physically independent, syntrophically connected hosts and symbionts. Models on the emergence of the initial ectosymbiosis through the evolution of attachment receptors (i.e., case 1 to 2 in Fig. 1) are discussed there.
- Stability of syntrophic interactions, assuming mutually beneficial metabolite-exchange [7]. There the model assumes faster metabolite dynamics compared to population dynamics (unlike the setup in this study).

Note that the entire analysis in this paper relied on the assumption that mechanisms to attach symbionts onto the host surface (e.g., membrane receptors) evolved earlier than the host developed abilities to fold its plasma membrane. The assumption was based on the idea that bioenergetics involved in modifying the host cell shape seem more expensive to that of expressing surface receptors. Rather than attempting to justify the exact order of appearance of these innovations, we focused on the factors behind the transition promoting better microbial integration. However, we emphasize that the chronology of events leading to integration in the context of endosymbiotic origin of mitochondria (or the eukaryotic ancestor) needs further investigation.

The larger contact surface area of the hosts with the ectosymbionts due to the membrane folds proves to be beneficial to the hosts in protecting them from the growth-inhibiting effect of self-produced and/or external metabolites. We observed that the membrane-folding host phenotype can emerge if the effective surface coverage is large enough to compensate for the effects of metabolic self-inhibition, despite its additional structural and energetic costs. Note that we specifically considered and modelled the stress that inhibits host growth to be purely metabolically induced. However, there could be other biological stresses acting on the host such as predators or phages (that are host-independent and require a different modelling approach), where the host might as well benefit from surface-covering symbionts [39]. Our results suggest that a metabolically induced toxic stress from the environment could be an incentive to form better-integrated microbial associations, purely from a unilateral metabolic relationship. This also does not require mechanisms like direct phagocytotic inclusion that could be prohibitively energy-demanding [40] or not [41], but is certainly lacking among archaea so far.

Eukaryotes often form metabolic interactions, and they can easily capture and retain other species via phagocytosis. The mitochondrial ancestor, on the other hand, is assumed to have been integrated by an archaeon, likely before phagocytosis was evolved in the ancestral host lineage [42]. According to syntrophic hypotheses on the origin of mitochondria, the initial interaction between the ancestral host and symbiont was mutually beneficial metabolic syntrophy [18,43–46]. Yet, prokaryotes, while many (or most) are syntrophic, mostly lack mechanisms to physically capture their partners [40,42]. Archaea, while having a dynamical cytoskeleton and endocytic components largely evolutionarily continuous with eukaryotic counterparts [47–49], are not known yet to possess endocytic (let alone phagocytotic) capabilities. There are known bacterial species capable of endocytosis [41,50], but it is unknown whether they can capture and retain living cells and whether the engulfed cell can survive long enough to provide metabolic help to be considered anything like an endosymbiont.

For evolution to act on the partnership, it is required that the two species reappear better than random over generations. Given no phagocytotic inclusion, prokaryotes had to evolve other means to retain their cooperative partners and ensure their stable vertical inheritance (unless they coexisted in the same habitat for a very long time, potentially without the interference of third parties and environmental perturbations). There are extant examples of obligate metabolic, syntrophic partnerships among prokaryotes (see, e.g., *Chlorochromatium* [13,21,51,52]), but we have no information either on their original interaction or on the evolutionary trajectory of their coevolution.

Without explicit internalization via phagocytosis, there are at least three potential ways of improving the contact with a partner (so far neither of them is known to have enabled endosymbiosis either in prokaryotes or in eukaryotes):

- Invaginating the plasma membrane to trap the partner (as some pre-phagocytotic theories imagine, e.g., [24,27]); observed in protists housing ectosymbionts [19,53,54].
- Growing membrane protrusions (with cytoplasm within) to potentially wrap and entangle the partner (as the inside-out hypothesis imagines, e.g., [25,26,55]).
- Growing pili or other cell-surface appendages; though these do not involve membrane-restructuring like mechanisms of the previous points, this seems to be the extant way how microbes increase uptake and contact surface among each other to enhance, e.g., the exchange of electrons [56–58], metabolites, DNA [59,60], antimicrobial compounds [61] and potentially other molecules [58,62].

Note that topologically the above listed mechanisms are equivalent and we do not differentiate between them from a modelling point of view. However, from an evolutionary perspective, they may very well differ: pili do not seem to offer a mechanistic way for the host to bind the partners long enough to ensure their vertical inheritance, though there are pili involved in cell adhesion, e.g., during infection [61,63] which serves as an example of symbiont-initiated contact rather than host-initiated contact (theoretically explored in [28,38]). On the other hand, invaginations and protrusions of the host may, theoretically, trap partners physically more easily, even if only as ectosymbionts. Vertical inheritance of symbionts (better-than-random co-occurrence of partners) is necessary for the pair to be selected for as a unit of evolution. If there is a selective advantage for the pair, evolution is expected to select consortia with better trapping abilities.

Membrane invaginations presumably can evolve without a symbiont providing metabolites for the host. Feeding in *Uabimicrobium* starts with invagination [41]. Theoretically, the ancestral host may have formed pockets to capture and digest prey externally as a precursor of phagocytosis [24,27,64,65]. Also, microbes can increase their metabolically active surface in other ways too. Photosynthetic or other autotrophic species primarily reliant on membrane bioenergetics involve the extensive invaginations of the inner membrane (e.g., Planctomycetes, anammoxosomes, carboxysomes, ferrosomes, mitochondrial cristae, cyanobacteria, many other intracytoplasmic membranes [66], even in Archaea [67]). Internal invaginations have the means to compartmentalize the host metabolism and separate biofunctions to increase their efficiency and decrease interference. Membrane invaginations appear in epibiont-bearing protists [19] and bacteria [22], and also in Asgards, with external material trapped (but no epibionts) [68]. Alternatively, outward membrane folding may form protrusions, observed in Asgards [14,18,69].

Furthermore, we point out that the inward membrane folding is likely beneficial for the symbiont for multiple reasons. The one that we have modelled is the increased access to host-produced metabolites. The other would be a safe habitat where symbiont could avoid predators or harmful factors of the environment (we have not modelled these, focusing solely on the metabolic benefit). Captured ectosymbionts could presumably continue to reproduce asexually (or even sexually, as partners are always around), which could provide a continuous resupply of partner individuals.

Additionally, beneficial mutants of the symbiont would likely remain in the vicinity of the host, positively group-select the consortium (long-term association) even without perfect vertical inheritance of the symbiont.

Another alternative to membrane invaginations or protrusions in our context is the formation of external appendages or pili-like structures on the host. These can also potentially increase contact surface for inhibition reduction without compromising entire active membrane surface for autotrophy. As our results suggest, if such structures insist higher energy requirements, the cost-benefit trade-off gets subsequently shifted to prevent invasion by those phenotypes and favors the residing one (whatever they maybe). Arguably, membrane invaginations are feasible only when the partner sizes are different, while same-size partners tend to use pili-like structures. Additionally, as mentioned before, pili-like structures or appendages are unlikely to maintain long-term associations compared to invaginations. However, a differentiated analysis and comparison of both scenarios are beyond the scope of our model and require further investigation.

A different approach to reduce inhibition by external metabolites is ectosymbiosis with full coverage of the host by the symbionts. However, the membrane-bending host is trivially favored over full-scale occupancy of the host surface because of significantly reduced food uptake by the host in the latter scenario. But imagine a case where the food intake is not hampered by coverage; for instance, the host food is supplied by the symbionts and not acquired passively from the environment (bilateral cross-feeding). The presence of a partner in host proximity that can provide them with food and simultaneously reduce inhibition by consuming their self-inhibiting product could be positively selected for, being directly and mutually beneficial for both parties. There is no additional cost of reduced consumption because the food to the host is supplied directly by the ectosymbionts without any passive diffusion. In this scenario, membrane bending seems to be not as necessary as in our original case (recall that ability to maintain some free surface for metabolite inflow without fully compromising inhibition reduction was the advantage of membrane-bending). If we were to consider bilateral syntrophy, it seems trivial that such a mechanism is favored over membrane bending. Assuming that the symbiont gives something back to the host renders their interaction mutually beneficial. However, here we have deliberately modelled the case without any such additional or direct benefit.

We focused on the worst-case and more likely scenario of unilateral metabolic relationship, where reduction in self-inhibition acted as the incentive (indirect benefit) for the host to evolve mechanisms that can bring the partner closer. We see that the mutant host with membrane bending can evolve purely due to the benefit of its symbiont protecting it from its own inhibitive product. Any additional benefit would clearly make it easier for the mutant consortium to evolve. The outcomes are less trivial when the metabolic exchange is asymmetric (also compare [7] and [70]).

Cross-feeding (in fact, any metabolic exchange) also benefits from physical proximity of the partners as a result of reduced diffusion and privatization of metabolites. However, it is not trivial whether bilateral, mutual metabolic interaction (cyclical syntrophy) would benefit more from membrane-bending than from inhibition-reduction, but this is certainly something that could and should be modelled at least theoretically. Given the unlikeliness of immediate mutualistic partner specificity, it is reasonable to assume that bilateral cross-feeding between the host-symbiont pair might have evolved *after* the unilateral syntrophic symbiont was enfolded by the host in order to stabilize the pair even more. The unilateral syntrophic interactions, where the host produce is utilized by the symbiont and the symbiont does not return the favor explicitly in terms of beneficial substrates (as we have modelled), are much more prevalent compared to mutual cross-feeding that requires partner and metabolite specificity. Our result suggests that the host-symbiont metabolic interaction need not be mutualistic in terms of either party producing the required metabolite for the other.

Additionally, our result proving that the toxic metabolite need not be host- or symbiont-produced means that close physical ectosymbiosis could emerge not only because of unilateral cross-feeding (assumed in our previous works) but due to protective mutualism in a harsh environment. An early theory for mitochondrial origin assumed that the benefit of mitochondria for the host was their capacity to sink oxygen presumably toxic for an anaerobic host [71,72], more recently taken up by Imachi et al. [18]. The problem with this is that any such detoxifying benefit immediately disappears when the symbiont moves inside the host as then oxygen has to diffuse through the host. Moreover, the host is now believed to be at least oxygen-tolerant or micro-aerophilic in light of Asgards [45,73]. While it is unclear what other external metabolites could have been sunk by the mitochondrial ancestor, we may not exclude that ectosymbioses (mitochondrial or general) evolved to provide protection against harmful external metabolic factors. Admittedly, however, if the symbiont is not dependent on host product but on some readily available metabolite, it is harder to explain why selection favors a host-attached partnership. In both cases, we assume that it is the host that binds the partner, but in the latter case, there is no incentive at all for the symbiont to get closer to the host.

## Conclusions

Metabolically linked syntrophic microbial partners are extremely common, especially among prokaryotes, and some may even evolve mechanisms to physically enhance integration between parties. Yet the driving forces behind the phenomenon are not well-established. Additionally, such metabolic syntrophy is often assumed as the starting point of mitochondrial endosymbiosis, with no phagotrophy available. Here, we have tested the hypothesis that the capacity of a close symbiotic partner to alleviate the host’s self-inhibition by consuming its product could enable stronger physical integration. This incentive could manifest via the evolution of morphological adaptations in the form of membrane restructuring to facilitate inter-partner contact. We analyzed this transition using a mathematical model, demonstrating how metabolic interactions can complement emergence of traits enhancing extent of physical association. We showed that environmental metabolic stress could incentivize microbial partners already in ectosymbiosis to form more compact (i.e., physically better-integrated) symbiotic consortia. Our model theoretically supports the potential existence of intermediate stages between free-living partnership and full-internalization via engulfment necessary for progressive, syntrophy-based endosymbiosis.

## Methods

### 1. Resident ectosymbiotic consortium

Consider an already established ectosymbiotic consortium as a collectively replicating entity of a host and symbionts on its membrane surface (Fig. 2). The consortium linearly feeds on a resource W (of constant available concentration *w* in the habitat) at a constant rate (*k*_R_) and metabolizes the consumed substrate for cellular subsistence (*l*_R_) and replication (*z*_R_). Based on a population growth formulation in previous works [28,38], considering metabolite dependence, life history parameters and scaling to account for consumption to growth translation of biomass, the consortium population grows at the rate *r*_R_: = (*k*_R_*w* − *l*_R_)(ln 2)/*z*_R_ − *d*_R_, where *d*_R_ is the intrinsic death rate.

The consortium, after metabolism, generates a by-product (metabolic waste) U at a rate proportional to its uptake of W. A part of the metabolic produce is directly consumed by the *n*_R_ syntrophic ectosymbionts on the host surface at a constant rate (*e*_R_), while the residue is externalized into the environment of volume *Ω* at the rate *κ*_R_: = (1 − *n*_R_*e*_R_)*k*_R_*w*. In addition to the direct feeding from the hosts, the ectosymbionts in a consortium also consume the metabolite U dispersed and diluted in the local environment (indirect consumption rate *ξ*_R_: = *n*_R_*e*_R_/*Ω*, where habitat volume parameter *Ω* introduces a scaling to reflect uniform metabolite dispersion in whole habitat volume and resultant dilution). The host metabolite also decays linearly at a constant rate *δ*.

Consortium population growth is inhibited by metabolite U produced by the host conspecifics in the environment where total concentration of host metabolite is *u*. The metabolite-induced self-inhibition is represented using a non-linear function (comparable to that used in [31]): *u*/(*a*_R_ + *bu*), where 1/*b* denotes the maximal inhibition and *a*_R_/*b* is the half-saturation constant. Here, *a*_R_: = *a*/(1 − *n*_R_*s*_R_), where *a* correspond to the maximal value of the growth inhibition factor when the hosts are free-living without any ectosymbionts (*n*_R_ = 0) and *s*_R_ is the impact of contact surface per ectosymbiont in modifying the inhibition factor. The ectosymbiotic association with surface coverage reduces the growth inhibition on the hosts (0 < *n*_R_*s*_R_ < 1). When *c*_R_ is the resident consortium population density and *u* is the concentration of host metabolite in the habitat, the following dynamical system represents the metabolite-dependent population dynamics:

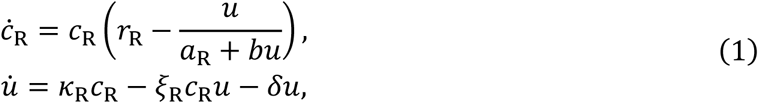

where all parameters are strictly positive. For the stability analysis of system (1), see Appendix A in Additional file 1.

### 2. Invasion by membrane-folding phenotype

Now consider the potential invasion of a mutant host phenotype capable of bending its membrane to potentially enfold the ectosymbionts, without crossing the plasma membrane. This partial engulfment increases the contact surface between ectosymbionts and their host compared to the resident ectosymbiotic consortium (*s*_M_ > *s*_R_) and can improve the chance of vertical inheritance of partners. The mutation is assumed to involve specific proteins and energy to bend the membrane inwards or outwards, thus incurring additional costs of subsistence and replication on the host (*l*_M_ > *l*_R_ and *z*_M_ > *z*_R_). The intrinsic death rate of the mutant is also considered higher than the resident to assume the worst-case scenario for the mutants (*d*_M_ > *d*_R_). Feeding of the mutant host is constrained by reduction in the free external surface (*k*_M_ < *k*_R_). The combined effect of these considerations on the life history parameters yields a lower intrinsic growth rate for the mutant host compared to the resident (*r*_M_ < *r*_R_). We assume the metabolite-specific inhibitory effects on both host phenotypes are identical. The only additional benefit the mutant host receives is that the effective growth inhibition by its own product U is reduced due to the increased contact surface with ectosymbionts (*a*_M_ > *a*_R_). The reduced exposure to the environment creates a geometric benefit to the mutants such that *a*_M_ = *a*/(1 − *n*_M_*s*_M_) > *a*_R_ = *a*/(1 − *n*_R_*s*_R_). Effectively, the mutant benefits from inhibition reduction while enduring an intrinsic growth rate inferior to the resident. Additionally, the symbionts help their host by lowering and protecting them from the toxic host metabolite in their vicinity. The ectosymbionts on the mutant host have the benefit of higher consumption of host metabolite U (*e*_M_ > *e*_R_ and *ξ*_M_ > *ξ*_R_). As a result, the host metabolite release is higher for the resident (*κ*_M_ < *κ*_R_). Recall the fundamental difference between the phenotypes is in the per-ectosymbiont contact surface area (*s*_M_ > *s*_R_). Thus, for easier comparison of phenotypes, the average ectosymbiont count on both host phenotypes is considered identical (*n*_R_ = *n*_M_ ≔ *n*). The dynamics of the two ectosymbiotic phenotypes and the external metabolite concentration, with all parameters strictly positive, can be represented as:

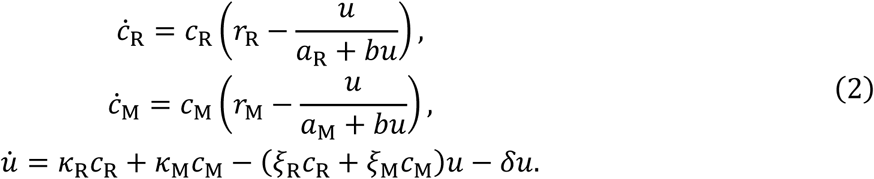

Results of stability analysis of system (2) are in Appendix B in Additional file 1.

### 3. Generalized model

Recall that, in the dimorphic model, we compared an already stable ectosymbiosis with a form of membrane folding (both without maximum surface coverage). Contrarily, here we relax our assumption of the stable ectosymbiotic consortium as the starting point of our analysis (i.e., as the resident) and explicitly demonstrate its stability against other possibilities of attachment without membrane folding. Additionally, to analyze the invasion by the membrane-folding mutant phenotype, we generalize the model to include multiple possible scenarios, i.e., non-ectosymbiotic and fully covered hosts considered simultaneously.

The different host associations or consortia vary in their capacity to accommodate symbionts and the extent to which they capture and retain each symbiont on their membrane. To include the several cases of Fig. 1, we consider the following host types: 1) non-folding and free-living (non-ectosymbiotic), 2) non-folding and ectosymbiotic without maximum surface coverage, 3) non-folding and ectosymbiotic with maximum surface coverage, 4) folding and free-living (non-ectosymbiotic), 5) folding and ectosymbiotic without maximum surface coverage and 6) folding and ectosymbiotic with maximum surface coverage. Let index *i* in subscript of parameters correspond to each of the respective listed host types. The ectosymbiont count in each case is as follows: *n*_1_ = *n*_4_ = 0 corresponding to the absence of ectosymbionts, *n*_3_ = *n*_6_ = *N* corresponding to maximum host surface coverage by *N* ectosymbionts and *n*_2_ = *n*_5_ = *n* (0 < *n* < *N*). The surface contact per ectosymbiont varies only between the non-folding and folding types: *s*_1_ = *s*_2_ = *s*_3_ = *s*_R_and *s*_4_ = *s*_5_ = *s*_6_ = *s*_M_ (as in the dimorphic model). The above considerations result in the following relations for the intrinsic growth rate, growth inhibition factor, metabolite release rate and indirect consumption rate, respectively: *r*_1_ > *r*_4_ > *r*_2_ > *r*_5_ > *r*_3_ > *r*_6_, *a*_1_ = *a*_4_ < *a*_2_ < *a*_5_ < *a*_3_ < *a*_6_, *κ*_1_ = *κ*_4_ > *κ*_2_ > *κ*_5_ > *κ*_3_ > *κ*_6_ and *ξ*_1_ = *ξ*_4_ < *ξ*_2_ < *ξ*_5_ < *ξ*_3_ < *ξ*_6_. Effectively, the distinct definitions lead to varied benefits of inhibition reduction and costs of growth reduction across the host types.

However, considering the different cases of attachment and membrane folding traits in the same ecological dynamics contradicts the fact that mutation occurs rarely and the fundamental assumption of only one mutation in an ecological timeframe. Note that our hypothesis also considers that attachment evolved before membrane folding. (We assume this to focus on the ecological driver of transition from typical ectosymbiosis to enhanced integration, without concern about the chronology of these events.) Hence, we consider the above six cases in two separate sets (or subsystems). Presuming attachment prior to folding, the ecological dynamics between cases 1 to 3 stabilize before the membrane folding mutant emerges. The membrane folding mutant invades the stable state of the resident ecological dynamics of attachment trait. Hence, the ecological dynamics of membrane folding trait should consider the stable ectosymbiotic consortium (that dominates the subsystem with attachment) along with cases 4 to 6. The dynamics of the host types in the two separate subsystems, based on the dimorphic model of two phenotypes, can be generalized as:

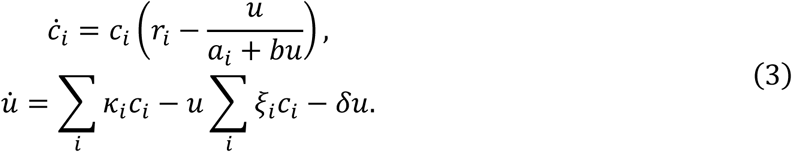

Appendix C in Additional file 1 contains the stability analysis of the general system (3).

The general dynamics for any mutant of trait value *θ* can now be defined as:

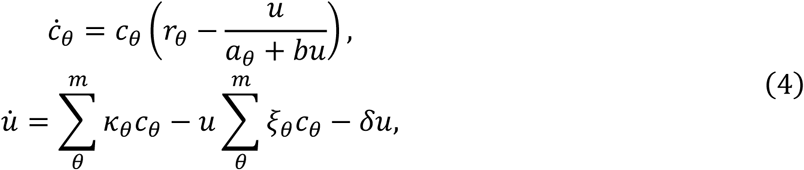

where *r_θ_* = (*k_θ_w* − *l_θ_*)(ln 2)/*z_θ_* − *d_θ_*, *a_θ_* = *a*/(1 − *ns_θ_*), *κ_θ_* = (1 − *ne_θ_*)*k_θ_w* and *ξ_θ_* = *ne_θ_*/*Ω* by generalizing terms introduced in Methods 1, and *m* is the actual number of distinct trait values (ecotypes) within the community. The simulation of the trait-dependent dynamical system (4) is initialized with a uniform host population with resident trait for which *s*_R_ = 0.2, *k*_R_ = 0.4, *l*_R_ = 5, *z*_R_ = 4, *d*_R_ = 0.33 and *e*_R_ = 0.2 as defined in Table 2 and Methods 1, and all initial densities in the dynamics are set to 0.01. At each timestep of *t* = 20, a new mutant is generated randomly by perturbing the trait value such that in generation *g* + 1, *θ_g_*_+1_ = *θ_g_* + *N*(0, *σ*), where the normal distribution has mean zero and standard deviation *σ* = 0.01. Mutants invade the population with initial density 0.01. During the simulation, any population density lower than 10^−5^ is considered extinct and is removed from the community. For more details on the simulation analyzing the trait evolution, see Appendix D and Fig. D1 in Additional file 1.

## Acknowledgments

NK is thankful to Gergely Röst for helpful discussions.

## Authors’ contributions

**NK**: Conceptualization, Methodology, Visualization, Formal analysis, Validation, Writing – original draft and Writing – review & editing; **JG**: Conceptualization, Methodology, Supervision, Writing – original draft and Writing – review & editing; **ÁK**: Conceptualization and Writing – review & editing; **MB**: Formal analysis and Writing – review & editing; **IZ**: Conceptualization, Methodology, Visualization, Supervision, Writing – original draft and Writing – review & editing. All authors read and approved the final manuscript.

