## Additional file 1: Appendix for "Metabolic inhibition can enable compact physical integration of microbial partners"

### Appendix A. Stability analysis of the monomorphic system

System (1) corresponding to the monomorphic dynamics in the main text has a unique interior fixed point $(c_{\mathrm{R}}^{*}, u^{*}) = \left(\frac{a_{\mathrm{R}} r_{\mathrm{R}} \delta}{\kappa_{\mathrm{R}}(1-b r_{\mathrm{R}})-a_{\mathrm{R}} r_{\mathrm{R}} \xi_{\mathrm{R}}}, \frac{a_{\mathrm{R}} r_{\mathrm{R}}}{1-b r_{\mathrm{R}}}\right)$ that exists if and only if $r_{\mathrm{R}} < 1/b$ and $\kappa_{\mathrm{R}}/\xi_{\mathrm{R}} > a_{\mathrm{R}} r_{\mathrm{R}}/(1 - b r_{\mathrm{R}})$. The linear stability analysis of system (1) using Jacobian matrices shows that the trivial fixed point $(0, 0)$ is always unstable and the interior fixed point (given it exists) is always asymptotically stable. Additionally, for the function $f(c_{\mathrm{R}}, u) = 1/c_{\mathrm{R}}$, the divergence $\nabla.(f \dot{c}_{\mathrm{R}}, f \dot{u}) = \partial(f \dot{c}_{\mathrm{R}})/\partial c_{\mathrm{R}} + \partial(f \dot{u})/\partial u = -\xi_{\mathrm{R}} - \delta/c_{\mathrm{R}}$ is always strictly negative for all positive values of $c_{\mathrm{R}}$ and $u$, i.e., divergence does not have sign change in the positive real plane. Thus, the Bendixson-Dulac criterion [1] suggests the system does not have any closed orbits in the positive real plane and the locally stable unique interior fixed point is also globally stable.

### Appendix B. Stability analysis of the dimorphic system

The ecologically feasible and non-trivial fixed points of the resident-mutant extended system (2) in the main text are $E_{\mathrm{R}} = (c_{\mathrm{R}}^{*}, 0, u_{\mathrm{R}}^{*}) = \left(\frac{a_{\mathrm{R}} r_{\mathrm{R}} \delta}{\kappa_{\mathrm{R}}(1-b r_{\mathrm{R}})-a_{\mathrm{R}} r_{\mathrm{R}} \xi_{\mathrm{R}}}, 0, \frac{a_{\mathrm{R}} r_{\mathrm{R}}}{1-b r_{\mathrm{R}}}\right)$ and $E_{\mathrm{M}} = (0, c_{\mathrm{M}}^{*}, u_{\mathrm{M}}^{*}) = \left(0, \frac{a_{\mathrm{M}} r_{\mathrm{M}} \delta}{\kappa_{\mathrm{M}}(1-b r_{\mathrm{M}})-a_{\mathrm{M}} r_{\mathrm{M}} \xi_{\mathrm{M}}}, \frac{a_{\mathrm{M}} r_{\mathrm{M}}}{1-b r_{\mathrm{M}}}\right)$. The existence and local stability conditions of the fixed points based on linearization are summarized in Table 1. Mutant invasion condition (see Fig. 4 in main text), that gives the stability of the mutant-only equilibrium $E_{\mathrm{M}}$, is:

$$\frac{r_{\mathrm{M}} a_{\mathrm{M}}}{1 - r_{\mathrm{M}} b} > \frac{r_{\mathrm{R}} a_{\mathrm{R}}}{1 - r_{\mathrm{R}} b}. \qquad (\mathrm{B}.1)$$

If the inequality is not satisfied, the resident fixed point $E_{\mathrm{R}}$ is stable (i.e., the resident is resistant to mutant invasion). If neither fixed point is stable (i.e., if neither pair of the existence and stability criteria in Table 1 is fully satisfied), the population densities blow up and do not stabilize. Rearranging condition $(\mathrm{B}.1)$, observe that the costs (reflected in $r_{\mathrm{M}}$) to the focal phenotype (i.e., the invading mutant) must be such that growth rate satisfies the following inequality:

$$r_{\mathrm{M}} > \frac{r_{\mathrm{R}} a_{\mathrm{R}}}{a_{\mathrm{M}} + a_{\mathrm{R}} b r_{\mathrm{R}} - a_{\mathrm{M}} b r_{\mathrm{R}}}. \qquad (\mathrm{B.2})$$

Note that the local stability analysis of the fixed point is applicable only when the mutant is introduced in rare densities. We now check if, regardless of the initial condition, the dimorphic dynamics can be pushed to the steady state with only the mutant. We analyze if the mutant-only fixed point of system (2), i.e., $E_{\mathrm{M}}(0, c_{\mathrm{M}}^*, u_{\mathrm{M}}^*)$, is globally asymptotically stable using the Lyapunov approach [1,2]. Consider the following function which vanishes only at the fixed point $E_{\mathrm{M}}$ and is non-negative, continuously differentiable and radially unbounded in the non-negative octant:

$$V(c_{\mathrm{R}}, c_{\mathrm{M}}, u) = \lambda_1 c_{\mathrm{R}} + \lambda_2 \int_{c_{\mathrm{M}}^*}^{c_{\mathrm{M}}} 1 - \frac{c_{\mathrm{M}}^*}{x}\, dx + \lambda_3 \int_{u_{\mathrm{M}}^*}^{u} 1 - \frac{a_{\mathrm{M}} + b u_{\mathrm{M}}^*}{a_{\mathrm{M}} + b y}\, dy, \qquad (\mathrm{B.3})$$

where $\lambda_1, \lambda_2, \lambda_3 > 0$. We follow this standard Lyapunov structure typically used to analyze global behavior of predator-prey (Lotka-Volterra dynamics) and resource-based chemostat models [3–8]. For further calculations to prove global stability in system (2), we consider similar approaches used for a chemostat-based model [8]. We can show that, if the mutant invasion condition ($\mathrm{B.1}$, or equivalently $\mathrm{B.2}$) is satisfied, there exist constants $\lambda_1$, $\lambda_2$ and $\lambda_3$, such that the time derivative of function in equation $(\mathrm{B.3})$ $\dot{V}(c_{\mathrm{R}}, c_{\mathrm{M}}, u) \le 0$ for all feasible values of $c_{\mathrm{R}}$, $c_{\mathrm{M}}$ and $u$. Here the equality is obtained if and only if $u = u_{\mathrm{M}}^*$ and $c_{\mathrm{R}} = 0$ (which also gives $c_{\mathrm{M}} = c_{\mathrm{M}}^*$ uniquely from system (2)). By the LaSalle's invariance principle [2], global asymptotic stability of the mutant-only equilibrium $E_{\mathrm{M}}(0, c_{\mathrm{M}}^*, u_{\mathrm{M}}^*)$ follows.

### Appendix C. Stability analysis of the generalized system

For the general system (3) with multiple host types, the fixed point $E_i = (0, \ldots, c_i^*, u_i^*) = \left(0, \ldots, \frac{a_i r_i \delta}{\kappa_i (1 - b r_i) - a_i \xi_i r_i}, \frac{r_i a_i}{1 - r_i b}\right)$ exists if and only if $r_i < 1/b$ and $\kappa_i/\xi_i > r_i a_i/(1 - r_i b)$. Note that only boundary equilibria (each with at most one host type) exist; no two host types can coexist. The condition for type $i$ to dominate all other types in its concerned subsystem, i.e., exclusive stability of $E_i$ within a subsystem, is

$$\frac{r_i a_i}{1 - r_i b} > \frac{r_q a_q}{1 - r_q b} \; \forall q \neq i, \qquad (C.1)$$

where index $q$ corresponds to all types other than the focal $i$-th type, and $r_i$ and $a_i$ respectively denote the intrinsic growth rate and growth inhibition parameter of the $i$-th type. For the subsystem with attachment, $i = 1, 2, 3$; here, case 2 is likely to dominate owing to the balance between $r_2$ and $a_2$. Hence, for the subsequent subsystem with folding, $i = 2, 4, 5, 6$ and stability here is conditional based on the trade-off as in the invasion condition. $i = 2$ and $i = 5$ in the generalized case respectively correspond to the resident and mutant phenotypes described in the dimorphic model (denoted using subscripts *R* and *M* in system (2) and condition $(\text{B}.1)$). Condition $(\text{C}.1)$ shows that the host phenotype that can effectively optimize $r_i a_i/(1 - r_i b)$ would dominate as any mutant host with properties meeting this condition can invade and stabilize. Evidently, this can be the case for high growth inhibition factor $(a_i)$ achieved by phenotypes with increased contact surface. Hence, if sufficient time is provided for invasion, the population is always expected to be monomorphic and with traits that improve contact surface.

### Appendix D. Trait evolution and invasion analysis

Now let $\theta \in [0, 1]$ be a variable trait that defines the quantities (or parameters) that differ among different phenotypes of the consortial host, where $\theta = 0$ corresponds to the resident trait. Parameters are in a trade-off (a beneficial increase in one has a disadvantage in other(s)) which is defined by their dependence on the trait value (Fig. D1(A)). The contact surface of the host with their ectosymbionts $(s_\theta)$ increases with the trait value, where initial value (of the resident) $s_\text{R} = 0.2$ and $s_\theta$ saturates as the trait value approaches its upper limit. The host consumption rate $(k_\theta)$ decreases with the trait value, having an initial value $k_\text{R} = 0.4$. The cost of living $(l_\theta)$, cost of replication $(z_\theta)$ and intrinsic death rate $(d_\theta)$ increase monotonically with the trait, where initial values $l_\text{R} = 5$, $z_\text{R} = 4$ and $d_\text{R} = 0.33$, respectively. The consumption rate of ectosymbionts $(e_\theta)$ also increases along with the trait value and $e_\text{R} = 0.2$. We define the following functions to represent the trait-dependent properties of a mutant phenotype:

$$
\begin{aligned}
s_\theta &= s_{\mathrm{R}} + \frac{\tanh 2\theta}{4}, \\
k_\theta &= k_{\mathrm{R}} - \frac{\tanh 2\theta}{10}, \\
d_\theta &= d_{\mathrm{R}} + \frac{\exp\theta - 1}{10}, \\
l_\theta &= l_{\mathrm{R}} + \exp\theta - 1, \\
z_\theta &= z_{\mathrm{R}} + \exp\theta - 1, \\
e_\theta &= e_{\mathrm{R}} + \frac{\exp\theta - 1}{10}.
\end{aligned}
\tag{D.1}
$$

The above functions are arbitrarily chosen such that they are compatible with the characteristics and behavior of the respective model parameters (see Fig. D1(A)). Based on the above considerations, the benefit of inhibition reduction to the hosts increases with the trait, while growth rate decreases as the worst-case scenario (Fig. D1(B)). The parameter values of the mutant phenotype used for the numerical analysis of the dimorphic model (Fig. 3 in main text) correspond to approximation of the values for $\theta = 0.6$ in equation $(\mathrm{D.1})$.

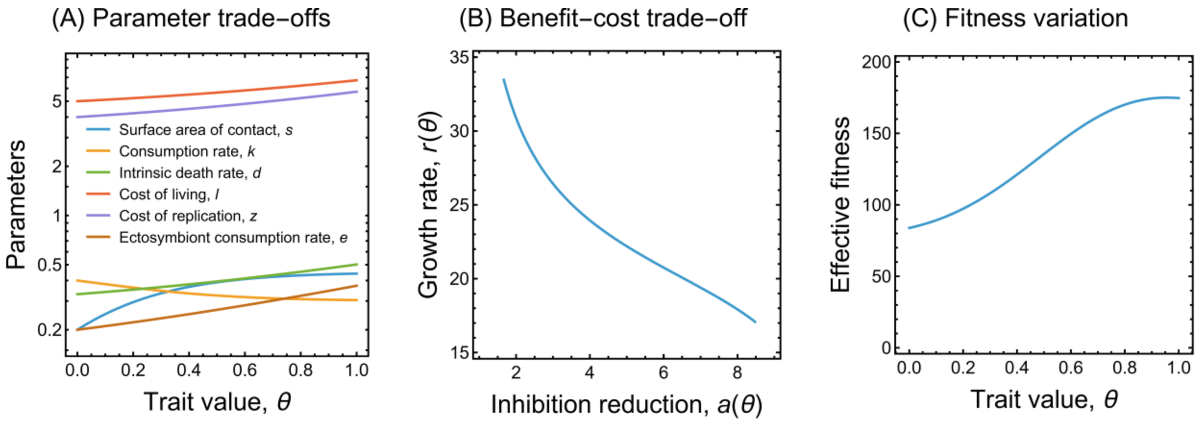


***Fig. D1*** *Trait invasion analysis. (**A**) Trade-off between the model parameters depending on the trait value. (**B**) The trait-dependent trade-off between inhibition reduction (benefit) and lowered growth rate (cost). (**C**) The trait-dependent effective fitness function ($r_\theta a_\theta/(1 - r_\theta b)$) is maximized at the trait value $\theta \approx 0.96$.*

Defining and following trait-dependent parameters (as in equation $(\mathrm{D.1})$) satisfying the characteristics, trait evolution (dynamics of system (4) using the methodology described in Methods 3; also see, e.g., [9]) shows that the trait is driven toward better physical integration (Fig. 5 in main text). The trait tends to fixate at the value that maximizes the function $r_\theta a_\theta/(1 - r_\theta b)$ (Fig. D1(C)).

### Appendix E. Competitive systems without self-inhibition

Here we check the possibility of membrane folding evolving without the additional stress of metabolite-induced growth inhibition. Assume the host's metabolite has no impact on its population growth, i.e., there is no self-inhibition. The population densities are now limited only by intraspecies competition. The new monomorphic resident system with strictly positive parameters can be represented as:

$$\begin{aligned} \dot{c}_{\mathrm{R}} &= c_{\mathrm{R}}(r_{\mathrm{R}} - c_{\mathrm{R}}), \\ \dot{u} &= \kappa_{\mathrm{R}} c_{\mathrm{R}} - \xi_{\mathrm{R}} c_{\mathrm{R}} u - \delta u. \end{aligned} \tag{E.1}$$

The unique interior fixed point $\left(r_{\mathrm{R}}, \frac{r_{\mathrm{R}}\kappa_{\mathrm{R}}}{\delta + r_{\mathrm{R}}\xi_{\mathrm{R}}}\right)$ of system $(\mathrm{E.1})$ is always stable unconditionally, while the trivial fixed point is always unstable. Similarly, the extended dimorphic system with the mutant in a purely competitive environment (without self-inhibition) can be represented as:

$$\begin{aligned} \dot{c}_{\mathrm{R}} &= c_{\mathrm{R}}(r_{\mathrm{R}} - c_{\mathrm{R}} - c_{\mathrm{M}}), \\ \dot{c}_{\mathrm{M}} &= c_{\mathrm{M}}(r_{\mathrm{M}} - c_{\mathrm{R}} - c_{\mathrm{M}}), \\ \dot{u} &= \kappa_{\mathrm{R}} c_{\mathrm{R}} + \kappa_{\mathrm{M}} c_{\mathrm{M}} - (\xi_{\mathrm{R}} c_{\mathrm{R}} + \xi_{\mathrm{M}} c_{\mathrm{M}})u - \delta u. \end{aligned} \tag{E.2}$$

The non-trivial fixed points of system $(\mathrm{E.2})$ are: $\left(r_{\mathrm{R}}, 0, \frac{r_{\mathrm{R}}\kappa_{\mathrm{R}}}{\delta + r_{\mathrm{R}}\xi_{\mathrm{R}}}\right)$ and $\left(0, r_{\mathrm{M}}, \frac{r_{\mathrm{M}}\kappa_{\mathrm{M}}}{\delta + r_{\mathrm{M}}\xi_{\mathrm{M}}}\right)$. The resident-only fixed point is always stable (as $r_{\mathrm{R}} > r_{\mathrm{M}}$), whereas the mutant-only fixed point is always unstable. The phenotype with the highest intrinsic growth rate dominates in a purely competitive system, and this result is valid even in the generalized model.

### Appendix F. Periodically varying metabolite concentration

Instead of assuming that the host's growth is inhibited by its own metabolic product (self-inhibition), now we consider a toxic metabolite produced neither by the hosts nor by the symbionts (say metabolite V). Let the concentration of metabolite V in the environment ($v$) be variable with maximal value of $v_{\max}$ and change sinusoidally in the range $[0, v_{\max}]$. We check whether inhibition by this independent metabolite in the environment affects the dynamics differently compared to self-inhibition. We retain the density-dependent competition between the two host phenotypes to make sure the host populations do not blow up. The new dynamics, with the metabolite V concentration changing with respect to time $t$ and independent of host and symbiont densities, can be expressed as:

$$\begin{aligned}\dot{c}_{\mathrm{R}} &= c_{\mathrm{R}}\left(r_{\mathrm{R}} - c_{\mathrm{R}} - c_{\mathrm{M}} - \frac{(v_{\max}/2)(1+\sin t)}{a_{\mathrm{R}} + b(v_{\max}/2)(1+\sin t)}\right),\\ \dot{c}_{\mathrm{M}} &= c_{\mathrm{M}}\left(r_{\mathrm{M}} - c_{\mathrm{R}} - c_{\mathrm{M}} - \frac{(v_{\max}/2)(1+\sin t)}{a_{\mathrm{M}} + b(v_{\max}/2)(1+\sin t)}\right).\end{aligned} \tag{F.1}$$

System $(\mathrm{F.1})$ demonstrates two possibilities where each of the host phenotypes can exclusively exist (Fig. F1).

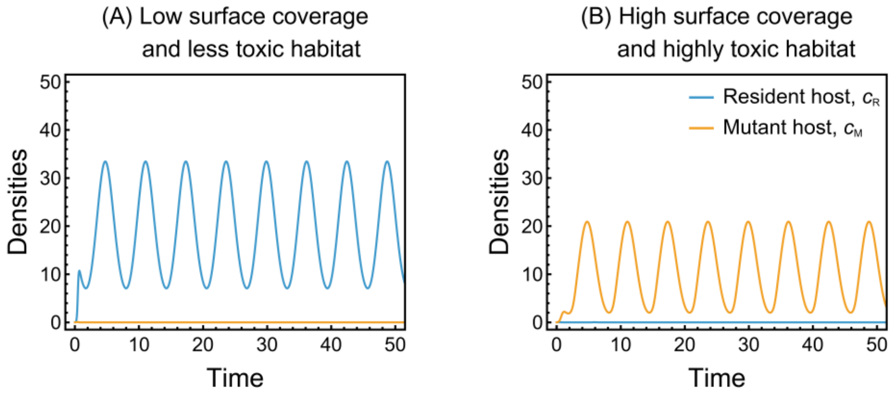


***Fig. F1*** *Population densities for periodically fluctuating metabolite concentration. Sinusoidally varying stress from the environment by a metabolite (not self-produced by the hosts or symbionts) can lead to either viable resident or mutant host population based on the growth inhibition factors of the phenotypes. For constant maximal concentration of metabolite V in the environment denoted $v_{max}$, low and high values of $v_{max}$ respectively correspond to less toxic and highly toxic habitats. Other parameters and their values are as in Table 2 in the main text. (**A**) $a_M/a_R = 1.2$ and $v_{max} = 60$ gives a non-zero resident population density. (**B**) $a_M/a_R = 4$ and $v_{max} = 120$ gives a non-zero mutant population density. The metabolic stress can create an incentive for better integration, even if the toxic metabolite is produced by neither party in the system, and the extent of surface coverage can influence the existence of the two host phenotypes.*
